# Multi-omics and deep learning provide a multifaceted view of cancer

**DOI:** 10.1101/2021.09.29.462364

**Authors:** Bora Uyar, Jonathan Ronen, Vedran Franke, Gaetano Gargiulo, Altuna Akalin

**Affiliations:** Bioinformatics and Omics Data Science Platform, Max Delbrück Center (MDC) for Molecular Medicine, The Berlin Institute for Medical Systems Biology, Hannoversche Str. 28, 10115 Berlin, Germany; Molecular Oncology, Max Delbrück Center (MDC) for Molecular Medicine, Robert-Rössle-Straße 10, 13125 Berlin, Germany

**Keywords:** pan-cancer, multi-omics, deep learning, variational auto-encoders, precision oncology, cancer subtypes, survival analysis, drug response

## Abstract

Cancer is a complex disease with a large financial and healthcare burden on society. One hallmark of the disease is the uncontrolled growth and proliferation of malignant cells. Unlike Mendelian diseases which may be explained by a few genomic loci, a deeper molecular and mechanistic understanding of the development of cancer is needed. Such an endeavor requires the integration of tens of thousands of molecular features across multiple layers of information encoded in the cells. In practical terms, this implies integration of multi omics information from the genome, transcriptome, epigenome, proteome, metabolome, and even micro-environmental factors such as the microbiome. Finding mechanistic insights and biomarkers in such a high dimensional space is a challenging task. Therefore, efficient machine learning techniques are needed to reduce the dimensionality of the data while simultaneously discovering complex but meaningful biomarkers. These markers then can lead to testable hypotheses in research and clinical applications. In this study, we applied advanced deep learning methods to uncover multi-omic fingerprints that are associated with a wide range of clinical and molecular features of tumor samples. Using these fingerprints, we can accurately classify different cancer types, and their subtypes. Non-linear multi-omic fingerprints can uncover clinical features associated with patient survival and response to treatment, ranging from chemotherapy to immunotherapy. In addition, multi-omic fingerprints may be deconvoluted into a meaningful subset of genes and genomic alterations to support clinically relevant decisions.

**Graphical Abstract:** 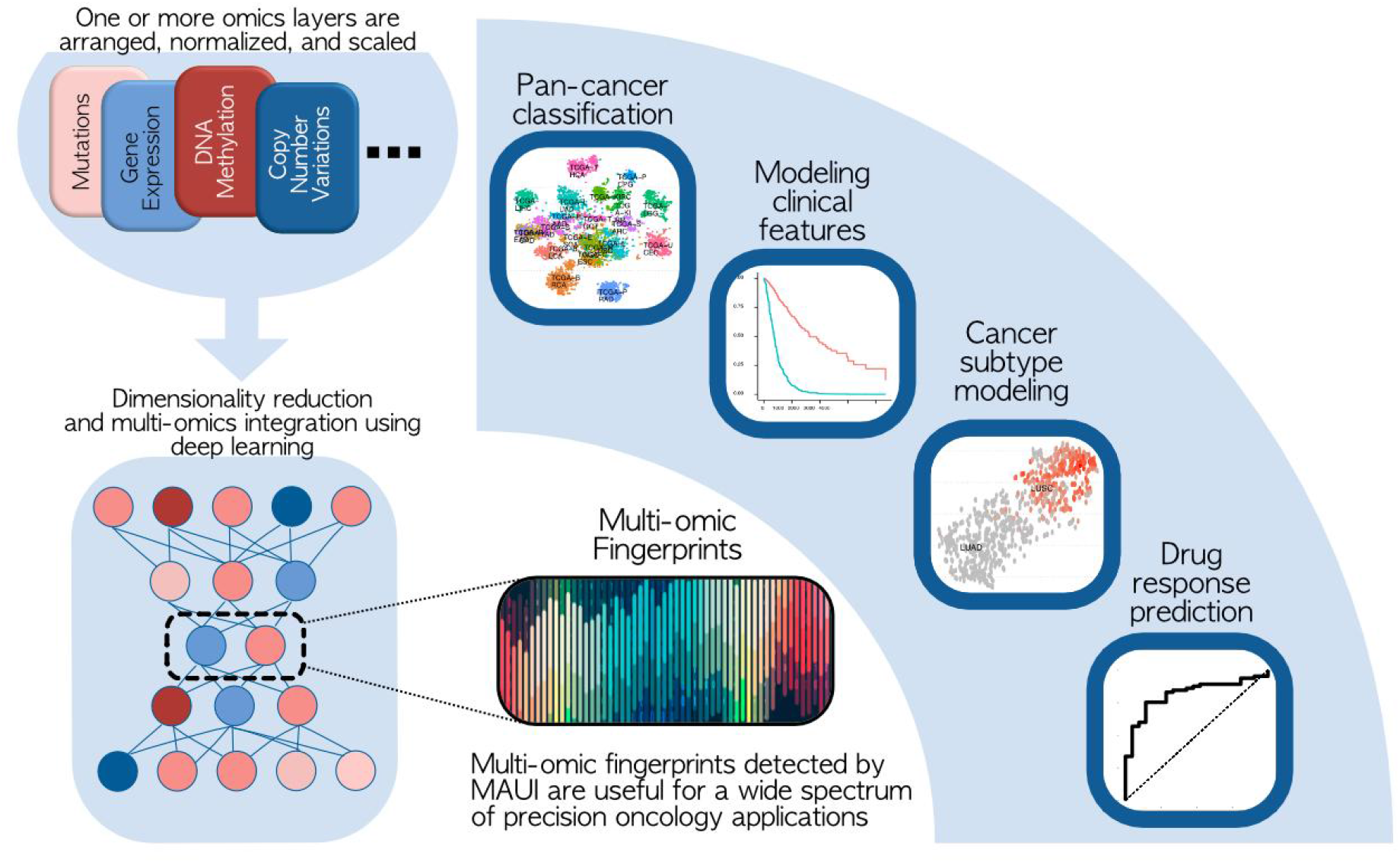

## Introduction

Cancer is a disease of the genome which is characterized by abnormal cell growth, invasive proliferation, and tissue dysfunction. It affected 19M people in 2020, and was the cause of 9.5M deaths in 2020 alone [1]. The underlying cause of the abnormal phenotype of cancer cells is, in most cases, acquired genetic defects that help cells circumvent safeguarding mechanisms. However, cancer cells must gain several cooperating hallmark features such as resisting cell death, avoiding immune system, tissue invasion, evading growth suppressors, and maintaining signaling for proliferation [2].

As opposed to rare diseases with Mendelian inheritance patterns, which are caused by few genomic variants that usually affect the protein sequence and structure, complex diseases such as cancer necessitate a deeper understanding of the combinatorial interplay between multiple cellular regulatory layers. This implies integration of data extracted from the omics layers such transcriptome, epigenome, proteome, genome, metabolome, and microbiome [3]. Genome-informed diagnostics, for instance, through the detection of disease-causing variants via exome sequencing, is already available in the clinic [4,5]. However, it is impossible to capture the complexity of most cancer types through the characterization of a few genomic markers. Multi-omics profiling of patients is a promising step towards such understanding, not only for cancer [6,7], but also for other complex diseases such as cardiovascular [8] or neurological diseases [9]. Proof-of-concept studies have already demonstrated the value of multi-omics profiling of patients for health monitoring [10] and treatment decision making [11]. Also underway are recent efforts of longitudinal clinical studies of cancer [12] comparing the impact of multi-omics guided clinical decision making in comparison to standard of care. In fact, large international consortia recognised the need for multi-omics and produced multi-omic databases such as The Cancer Genome Atlas (TCGA), Cancer Cell Line Encyclopedia (CCLE) [13], TracerX [14] for deeper molecular profiling of the tumors and disease models.

While multi-omics profiling shows promise for cancer research and precision oncology, due to the high-throughput nature of the generated data, the integration of multi-omics datasets is a challenging procedure [15]. The inherent high dimensionality of multiomics datasets, combined with the heterogeneity of the collected data necessitates the application of specialized machine learning and deep learning methods. To this end, various multi-omics integration methods have been developed [15,16]. There are multiple strategies of integrating multi-omics data with regards to the order in which layers get integrated. While some earlier multi-omics integration studies have opted for a sequential integration approach, where each omics layer is analysed one after the other, more recently developed sophisticated methods have enabled *late*, and *joint* integration of different omics layers. With a late integration strategy, each omic data type is analyzed separately, and the results are integrated. Late integration schemes [17,18] excel at capturing patterns which are reproducible between the omics data types, but can be blind to cross-modality patterns. An alternative strategy is joint integration and dimensionality reduction, whereby the different data modalities are jointly analysed from the start [16,19–24]. Such joint integration and dimension reduction methods construct a common latent space representation from the different omics data types. The latent space representation is constructed in a way that it reproduces important patterns from the different omics types with a much lower dimensionality than the combined multi-omics feature space. The dimensions of the latent space are called latent factors, and each latent factor is a combination of multi-omic input features. Latent factors therefore capture the complex interactions between the multi-omic variables.

An important factor to take into consideration when modeling complex diseases such as cancer is that integration methods need to capture non-linear associations both within and between the different omics layers. Current methods for joint dimension reduction are not designed to address this task. To remedy this problem, we and others have developed deep neural network based approaches. The most commonly used deep neural network architectures for latent factor modeling of multi-omics datasets are autoencoders. Different variants of auto-encoder architectures have been used for the analysis of multi-omics datasets [23,25,26]. However, these studies have demonstrated the usefulness of the methods on only selected cancer types and have not extensively and comprehensively tested their methods on a variety of use cases.

In this study, we demonstrate the utility of our deep learning framework MAUI, which is a stacked beta-variational auto-encoder [23], for modeling clinical and molecular features of tumor samples across a spectrum of cancer types. The high dimensional multi-omic feature datasets are first reduced via MAUI into low dimensional latent factors. In order to distinguish the nonlinear and combinatorial nature of the MAUI-based latent factors from the latent factors generated by linear modelling approaches, we refer to MAUI latent factors as “multi-omic fingerprints”. We demonstrate that MAUI-based multi-omic fingerprints enable us to capture sources of biological variation among the tumor samples, which is reflected in the heterogeneous omics profiles across the selected 21 TCGA cohorts and can be used to learn features associated with survival outcomes. Moreover, we show that they can be used to predict and further characterise molecular subtypes of cancer, for instance, MSI status of pan-gastrointestinal cancers or histological subtypes for non-small-cell lung cancers. We also show that multi-omic fingerprints can be distilled further to simple decision trees for interpretability and practical purposes. Finally, we demonstrate how the multi-omic fingerprints can be informative in capturing features of response to the anti-PD-L1 immunotherapy in a metastatic urothelial cancer cohort, and how such fingerprints can be utilized to uncover well established biomarkers of chemotherapy resistance in a cohort of Glioblastoma Multiforme patients. Altogether, these use cases including benchmarking experiments against a select set of popular joint multi-omics integration tools demonstrate the clinically relevant applicability of deep learning-based multi-omic fingerprints for precision medicine approaches.

## Results

Multi-omic fingerprints computed for a given cohort of patient samples can be seen as abstract molecular patterns. They contain combinatorial contributions from various data modalities (as shown in Fig 1B). Here, we demonstrate that the fingerprints learned by MAUI contain clinically meaningful information that can be utilized in a variety of use cases for precision/personalized medical applications.

**Figure 1 :**
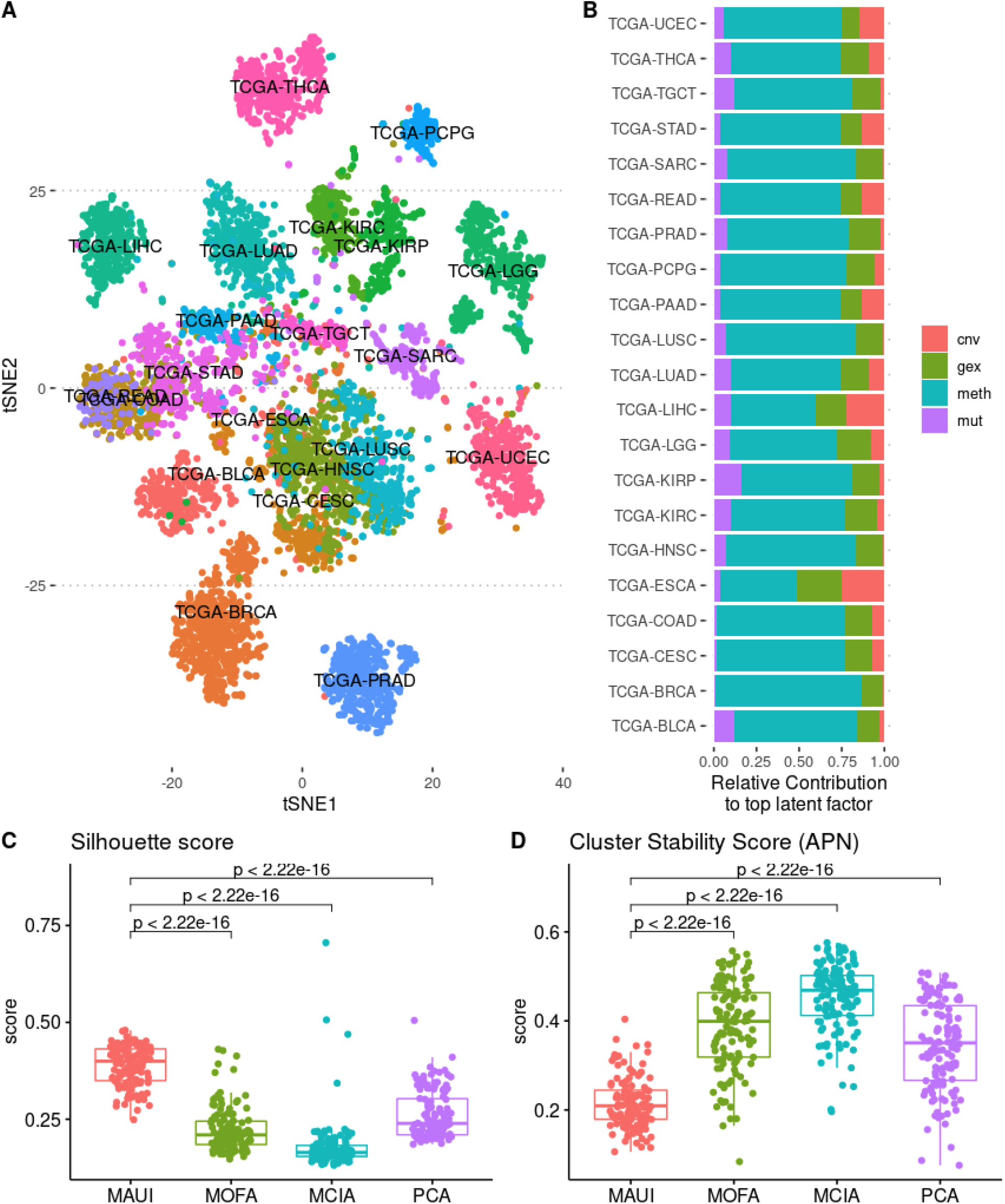
A) tSNE plot of MAUI multi-omics fingerprints (using top 5 fingerprints enriched per cancer type) learned from pan-cancer dataset B) Percentage of top 100 features by omics types contributing to top predictive MAUI fingerprint per cancer type. C, D) Comparison of unsupervised clustering performances based on cluster homogeneity (Silhouette) in C and cluster stability (Average Proportion of Non-Overlap (APN) in D. Each point represents a score for an individual clustering experiment in a total of 132 k-means clustering experiments (21 cancer types x for 6 different values of k from 3 to 8) for each tool.

### Multi-omic fingerprints can be used to precisely classify cancer types

We wanted to investigate whether the multi-omic fingerprints obtained using MAUI show any utility in distinguishing biologically distinct entities (tumors from different organs/tissues). We trained MAUI in a pan-cancer setting on 6775 samples from 21 cancer types (each cancer type containing at least 100 samples). We included only samples profiled with all four omics platforms (gene expression, methylation, copy number variation, and somatic mutations). The t-SNE [27] representation of the pan-cancer multi-omic fingerprints demonstrates that the samples mainly cluster by the cancer type and tissue of origin (Fig. 1A). In order to quantify how well MAUI separates samples from different cancer types, we trained an Elastic Net classifier on multi-omic fingerprints using 60% of the samples in a multi-class prediction. The classifier yielded a 92% mean-balanced accuracy on the testing dataset with a comparable performance to other multi-omics integration methods (Supp. Fig. 1A). The confusion matrix (Supp. Fig. 1B) of the predicted and the real cancer type labels of the samples shows that the majority of misclassified samples actually belong to the same organ system, for instance, the colon cancer (TCGA-COAD) and rectal cancer (TCGA-READ). Plotting the top multi-omic fingerprints predictive of each cancer type, we observed that multi-omic fingerprints show cancer-specific patterns with almost a one-to-one relationship (Supp. Fig. 1C). Moreover, top input features with the highest contribution to the cancer-predictive multi-omic fingerprints showed a variable distribution of data types, which was mostly dominated by DNA methylation (Fig 1B). This suggests that multi-omic fingerprints capture and summarize relevant multimodal information and integrate the most predictive value of individual omics layers into a single composite biomarker that can be used to precisely classify cancer types.

### Unsupervised clustering of samples using multi-omic fingerprints yields homogeneous and stable clusters

In the absence of sample-specific labels, the validity of clusters can be determined using statistical measures [28]. To evaluate the validity of the obtained sample clusters, we measured the Silhouette index (for cluster homogeneity) and the average proportion of non-overlap (APN) that measures cluster stability, separately for each TCGA cohort. Maximisation of within-cluster homogeneity together with inter-cluster heterogeneity serves the purpose of being able to detect cluster specific features, while cluster stability serves the purpose of the reproducibility of the biological conclusions drawn for each cluster. We clustered each cancer cohort with a selection of matrix factorization methods, and compared the performance of each tool against our multi-omic fingerprints obtained from MAUI. We observed that samples clustered using MAUI-based multi-omic fingerprints yield more homogeneous (Fig 1C) and more stable (Fig 1D) clusters in comparison to other methods.

### Multi-omic fingerprints capture meaningful clinical features

In a given cohort of patients with a common cancer diagnosis, the heterogeneity of the patient samples observed at the molecular level can be associated with the underlying biological variability of the samples. Sample-specific factors such as the tumor stage, localisation, histological subtype, or donor-specific factors such as the age, gender, and lifestyle habits (e.g. alcohol consumption or smoking) can all contribute to the heterogeneity observed in the omics profiles of each tumor sample. Latent factors derived from the multi-omics profiles should ideally reflect such known sources of variation. Moreover, these factors should also capture other unknown sources of variation, which when combined with known factors, should be useful in predictive tasks such as the survival outcomes of the donor or potential response of the donor to a specific treatment.

In order to demonstrate that multi-omic fingerprints can capture known sources of variation, we associated multi-omic fingerprints for each TCGA cohort to known clinical parameters. We looked for fingerprints that show differential distribution for a subgroup of samples compared to the rest of the samples (e.g. samples of female gender versus the rest). For 20 out of 21 cancers, we could find at least one multi-omic fingerprint associated with at least one basic clinical factor such as tumor stage, gender, histological type (with an adjusted p-value threshold of 0.05) (Fig 2A). For instance, the Lower Grade Glioma (LGG) is a heterogeneous set of diseases, which comprises strikingly different entities and MAUI effectively captures Oligodendrogliomas in fingerprint LF45 and histological grade III in LF45 (both p-value < 0.0001, Wilcoxon Rank sum Test; Fig 2B). Of note, LGG donors of male gender were enriched for MAUI fingerprint LF54 compared to females, which reflects the mild gender association observable in the metadata.

**Figure 2 :**
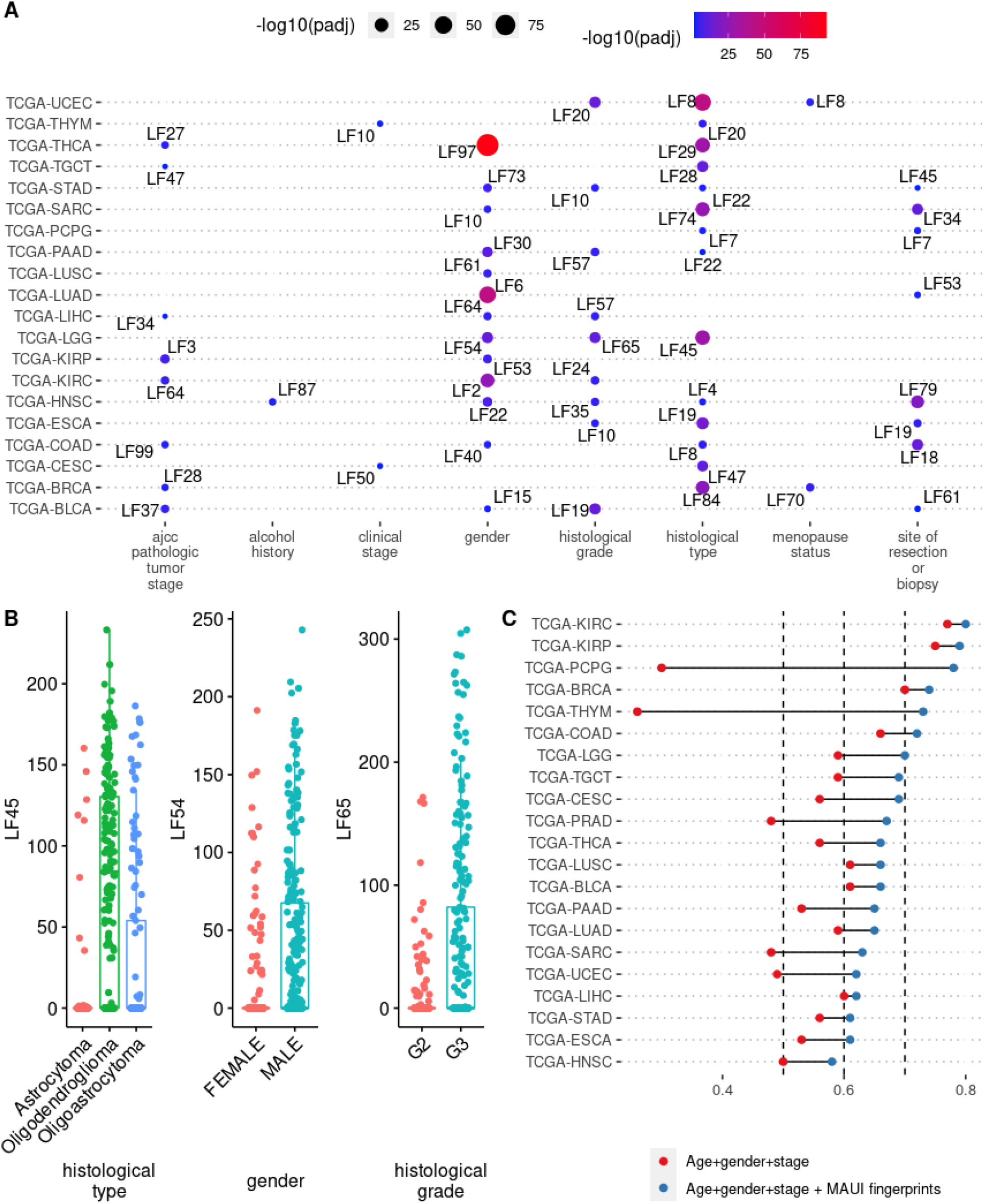
A) Top significant (padj < 0.05) clinical factor-associated multi-omic fingerprints per cancer type. B) MAUI fingerprints enriched for subgroups of different clinical features of TCGA-LGG samples. LF45 is enriched among samples histologically categorized as Oligodendrogliomas; LF54 is enriched among donors of male gender; LF65 is enriched among samples with histological grade of G3. C) Harrell’s C-index for progression-free survival prediction accuracy computed using only clinical factors (age + gender + tumor stage) or using clinical factors along with survival-predictive MAUI fingerprints

Secondly, we evaluated the power of the multi-omic fingerprints in combination with known clinical factors for predicting survival outcomes (progression-free interval). In comparison to the baseline survival outcome prediction accuracy (attained using only the clinical factors such as tumor stage, age, and gender), we demonstrated that adding the multi-omic fingerprints on top of such clinical factors improved the survival outcome predictions (Fig. 2B). We also demonstrated that our multi-omic fingerprints can be used along with basic clinical factors to prognostically stratify the patients (Supp. Fig. 2).

One further major source of variation between tumor samples is the composition of the tumor microenvironment. The tumor sample that is extracted for multi-omics profiling does not only contain cancer cells, but also other cell types such as the immune cells, fibroblasts, endothelial cells, and other non-cancerous cell types [29]. Tumor purity is a measure of the proportion of the cancer cells found in a tumor. The level of tumor purity for a given tumor sample can be a result of both the underlying biological mechanisms that lead to tumor growth and the technical protocol that is followed in surgically removing the tumor sample, which in either case has an impact on the multi-omics profiles obtained from samples [30]. Therefore, the tumor purity is an important confounding variable that needs to be estimated. Tumor purity estimates for the TCGA samples are already made available, which are based on either immunohistochemistry (IHC) or computational methods that are based on the analysis of somatic DNA alterations [31], gene expression [32], and leukocyte un-methylation [30]. These estimates are further assembled to obtain consensus tumor purity estimates (CPE) [30]. In order to evaluate the predictive power of MAUI multi-omic fingerprints for tumor purity estimates, we built Elastic Net regression models (using five-fold repeated cross-validation) on 60% of the samples for each TCGA cohort, where tumor purity estimates were available (14 TCGA cancer types out of the 21 studied cancer types). Then, we evaluated the models on the remaining 40% of the samples with respect to the correlation between the predicted tumor purity values and the reported tumor purity estimates by IHC or CPE. We obtained an average Pearson correlation coefficient of 0.30 across TCGA cohorts for IHC measurements (Supp. Fig 3A). For comparison, the average correlation between IHC values and estimates of other computational models are 0.31 for ABSOLUTE, 0.24 for ESTIMATE, and 0.21 for LUMP (Supp. Fig 3B). MAUI multi-omic fingerprints are also highly predictive of the consensus purity estimates (CPE) annotated for the TCGA samples for these cohorts with an average correlation of 0.74 across 14 TCGA cancer types (Supp. Fig. 3C). Example scatter plots of the MAUI predictions with respect to the annotated IHC and CPE values can be seen in Supp. Fig. 3D. These results suggest that MAUI multi-omic fingerprints can also reasonably capture the impact of non-cancerous cells in the tumor microenvironment on bulk multi-omics profiles, which are well expected confounding factors but also simple features discriminating otherwise similar biopsies.

### Multi-omic fingerprints can be used to classify and characterize molecular subtypes of cancers

#### MSI status in Pan-Gastrointestinal Cancers

Microsatellite Instability (MSI) is a molecular phenotype that is characterized by a high mutational load due to deficient DNA mismatch repair capacity [33]. A high level of MSI (MSI-H) is often observed in gastrointestinal cancers [34,35]. More importantly, MSI-H status has been proposed as a predictive biomarker for immune checkpoint blockade therapies [36–38]. Therefore, delineation of samples with MSI-H status is a clinically relevant objective for immunotherapy guidance. We trained MAUI on 867 pan-gastrointestinal samples using multi-omics features. To observe if MAUI LFs are predictive of MSI-H status of the gastrointestinal tumors we used the MSI status annotations from [35] as a dependent variable. The MSI status was quantified using a MSI-Mono-Dinucleotide Assay that examines mono/di-nucleotide repeat loci [35]. In comparison to other tools, MAUI multi-omic fingerprints were more strongly associated with the MSI status sub-groups (Fig. 3A). We also trained an Elastic Net [39] model with 5-fold cross-validation on 60% of the samples and reported the test set AUC values for the prediction accuracy. We observed that MAUI LFs are highly predictive of MSI-H status (AUC = 0.93) (Fig 3B). Later, we computed a reduced representation of the samples using a subset of learned MAUI multi-omic fingerprints, which are most predictive of MSI status. The tSNE plot of the samples colored by MSI status (Fig 3C) and the top predictive fingerprints (Fig 3D) demonstrates how well the specific fingerprint score correlates with the MSI-H status and can be used to distinguish samples with MSI-H from those with stable/low MSI status (Fig 3E). We inspected the top features that contribute to this specific multi-omic fingerprint and observed that top features predictive of MSI-H status are mostly populated by mutations and methylation sites (Fig 3F).

**Figure 3 :**
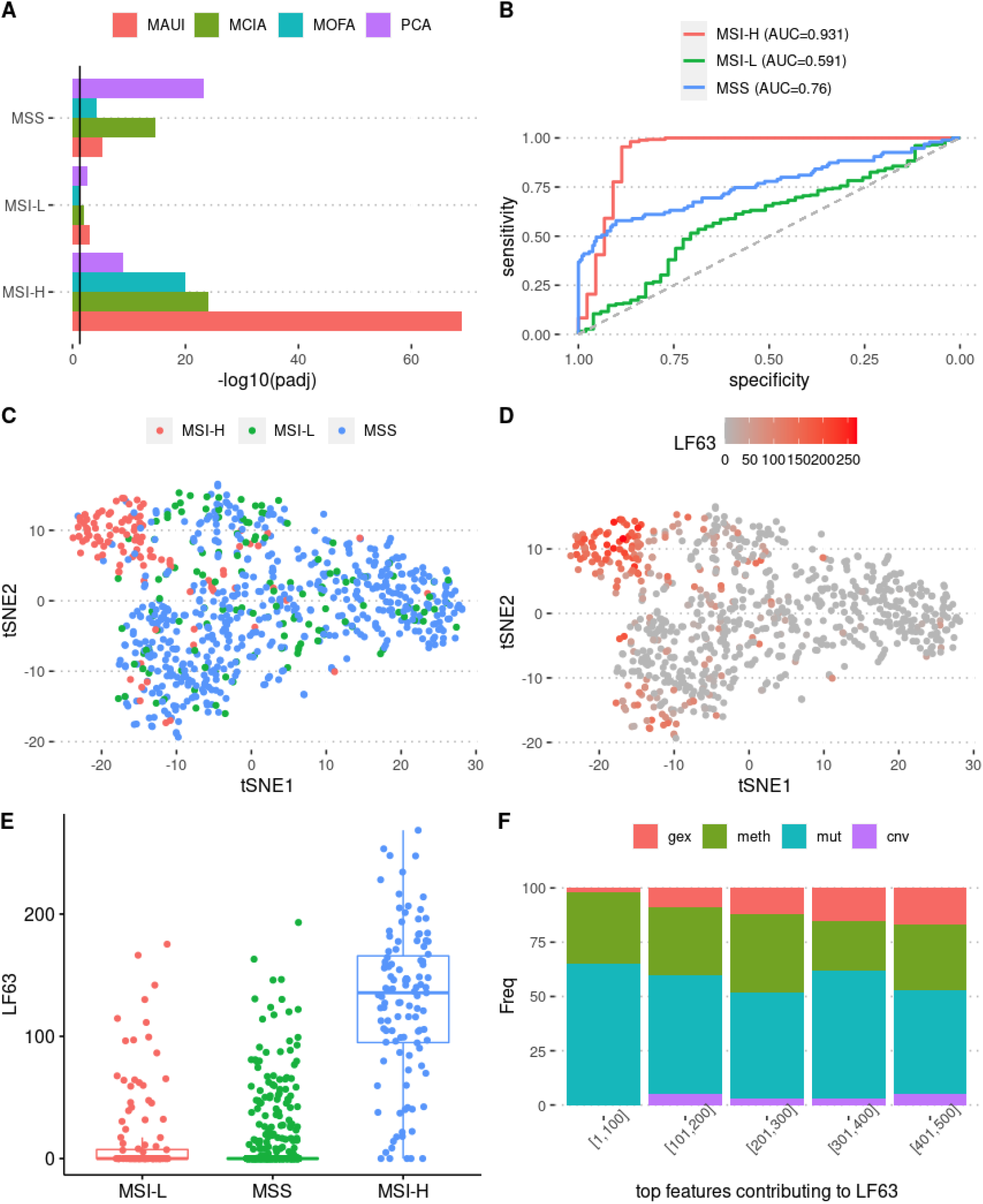
Latent factors predictive of MSI status (high: MSI-H, stable: MSI-S, low: MSI-L) in pan-gastrointestinal cancers A) Comparison of top latent factor per MSI status for each tool in terms of adjusted p-values reflecting the differences between MSI subgroups. B) ROC curves for prediction accuracy of MSI status using MAUI latent factors (multi-omic fingerprints) C) tSNE plots of samples generated using top MSI-predictive fingerprints (5 LF per MSI class) colored by MSI status D) tSNE plots of samples colored by MAUI fingerprint LF63, which is most predictive of MSI-H samples. E) Distribution of MAUI fingerprint LF63 across MSI subgroups in the pan-gastrointestinal cancer samples. F) Relative abundance of omics feature types among the top 500 features contributing to MAUI fingerprint LF63.

#### Classification of Non-Small-Cell Lung Cancers: LUAD vs LUSC

Lung Adenocarcinoma (LUAD) and Lung Squamous Cell Carcinoma (LUSC) are two histological subtypes of non-small-cell lung cancers (NSCLC). The treatment strategies for the NSCLC patients are mainly guided by the tumor stage rather than the histological subtypes. However, these subtypes display distinct pathway activities and cancer states [40–42]. Moreover, prognostic molecular predictors of disease recurrence are different between LUAD and LUSC subtypes [42]. Therefore, it is important to distinguish and characterize molecular features of these two types of non-small-cell lung cancers.

We trained MAUI on 800 NSCLC samples (441 LUAD samples, 359 LUSC samples) from the TCGA database. 5-fold cross-validated training of an Elastic Net model using the MAUI multi-omic fingerprints on the 60% of the dataset yielded a balanced AUC score of 0.98, a highly accurate classification performance on the test data (40% of the dataset) (Fig 4A-B). We confirmed that the activation scores of the best predictors for both LUAD (Fig 4C) and LUSC (Fig 4D) display the expected subtype specific activity.

**Figure 4 :**
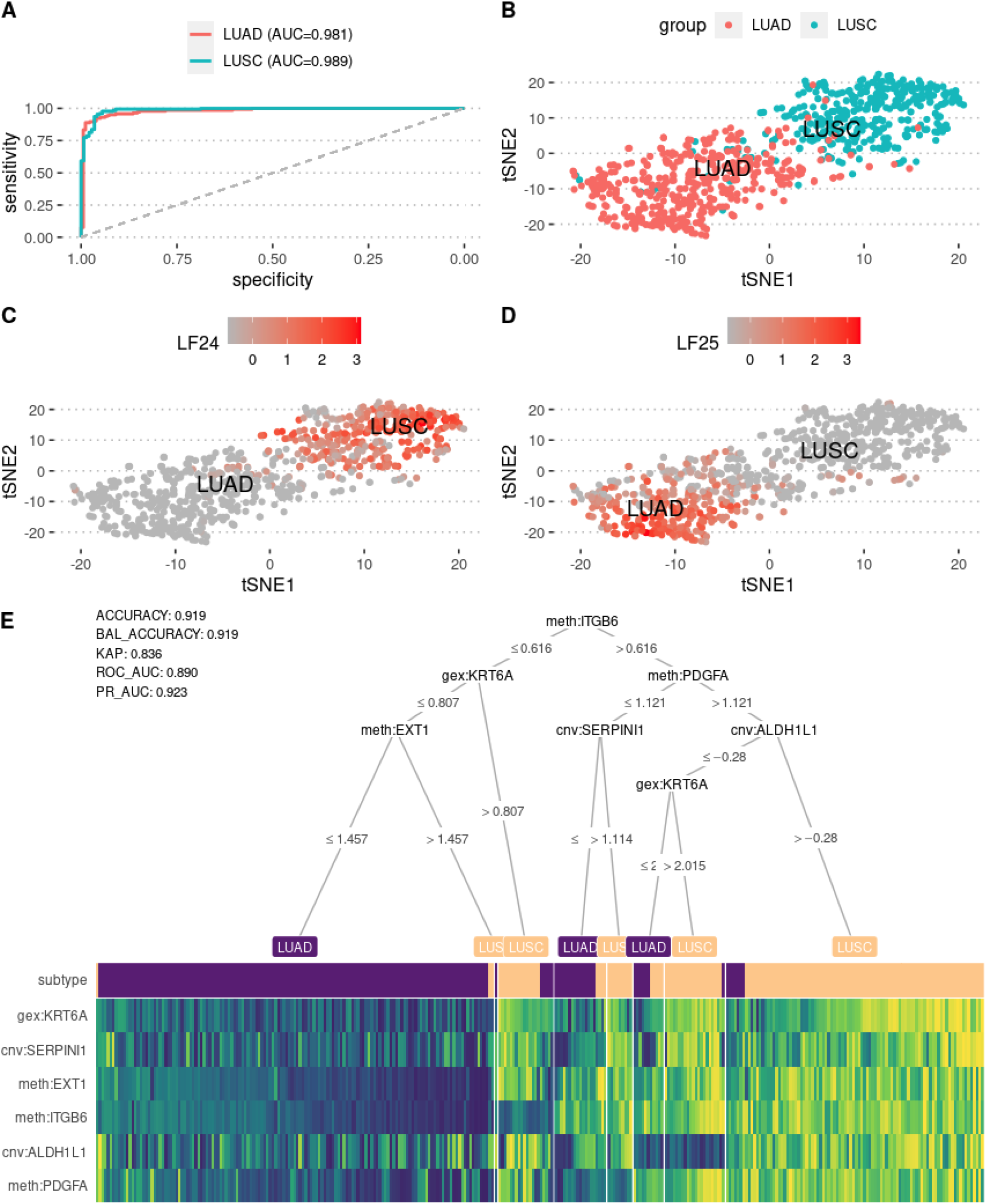
Classification of Non-Small-Cell Lung Adenocarcinoma samples using MAUI multi-omic fingerprints. A) ROC curves with AUC values on testing data (40% of the dataset) using an Elastic Net model with 5-fold cross-validation. B) t-SNE plot of MAUI fingerprints colored by cancer types. C) t-SNE plot of MAUI fingerprints colored by the top most-predictive fingerprint for TCGA-LUSC samples. D) t-SNE plot of MAUI fingerprints colored by the top most predictive fingerprint for TCGA-LUAD samples. E) Decision tree visualisation of top MAUI fingerprint-associated omics markers detected that can be used to distinguish LUSC and LUAD subtypes.

Although multi-omic fingerprints are highly predictive of the molecular subtypes, they might not be as clinically actionable as the input omics features. To increase the interpretability of the fingerprint-based classification, we extracted the top features associated with each of the fingerprints. We first computed top contributors (top 10 omics features per multi-omic fingerprint) to top subtype-specific fingerprints (top 5 per each subtype). We used these features to build an initial conditional inference tree. From this initial model, we picked the most important features and built a final decision tree that could be used as biomarkers of lung cancer subtyping (Fig. 4E). The single decision tree built in Fig. 4E, yielded a classification accuracy of 92%, however, when using these top 10 biomarkers in a more sophisticated classifier such as a Random Forest model, we obtained a classification accuracy of 98%, which is as good as what was achieved by the multi-omic fingerprints. This result suggests that multi-omic fingerprints can be used as exploratory tools to detect omics markers. Furthermore, the top detected biomarkers reflect the fact that MAUI fingerprints can capture interactions between different omics layers (copy number variation, methylation, and gene expression).

### Multi-omic fingerprints can be used to model treatment response

#### Predicting anti-PD-L1 treatment response in metastatic urothelial cancers

Despite the promise of precision oncology for guiding treatment decisions [43], only a minority of patients benefit from precision therapy based on the genetic information [44]. One of the most important breakthrough developments in precision medicine in the last decade, the cancer immunotherapy [45], could be made more efficient by recruiting the subset of patients that are more likely to respond to the specific kind of immunotherapy treatments. An example of such immunotherapy drugs is the anti-PD-L1 immune-checkpoint inhibitors, which has been recently studied in a cohort of patients (N=348) with metastatic urothelial cancers, who were profiled at whole transcriptome level before the start of the treatment and the patients were categorised based on their responses to the treatment [46]. We could detect MAUI fingerprints (in this case based on a single omics layer) associated with both responders (*p value* < 0.0001, Wilcoxon Ranksum Test) and non-responders to therapy (*p value <* 0.0001, Wilcoxon Ranksum Test) (Fig 5A). Moreover, we could detect MAUI fingerprints significantly associated with immune desert phenotype (*p value* < 0.0001, Wilcoxon Ranksum Test) (Fig 5B) and positively correlated with tumor mutation burden (*r* = 0.3) (Fig 5C), both of which are markers associated with immunotherapy responses that are proposed to be relevant for treatment decisions. Various MAUI fingerprints were found to be associated with response/non-response to therapy (Fig. 5D) and the fingerprint with the strongest association to response could be shown also as a strong predictor of progression-free survival outcomes (Fig 5E). Next, we built separate models to predict anti-PD-L1 responders using MAUI LFs alone or in combination with candidate biomarkers for immune checkpoint inhibitors (ICI) response such as tumor mutational burden (TMB), CD8+ T-cell effector gene signature, and pan-fibroblast TGF-beta response signature (F-TGBS) [46]. Whereas TMB has been linked to response to ICI in specific contexts [47], we show that MAUI fingerprints significantly add predictive power to established biomarkers, including but not limited to TMB (AUC = 0.82; Fig 5F). This suggests that MAUI may help to identify combinatorial biomarkers.

**Figure 5 :**
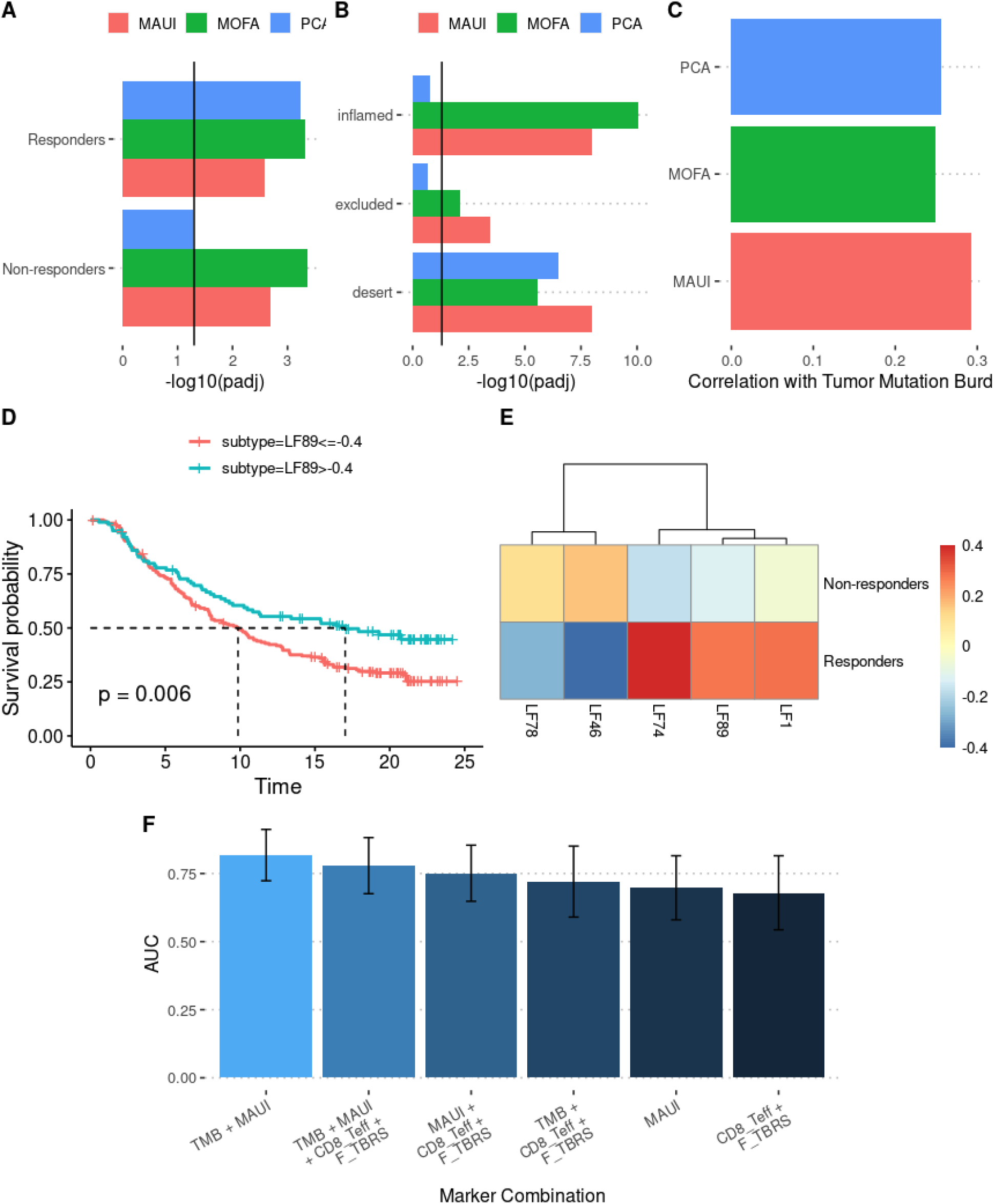
Predicting responders to anti-PD-L1 treatment in a cohort of Metastatic Urothelial Cancers (N=216) using MAUI. A) Comparison of top latent factors predictive of complete/partial response (CR/PR) to the anti-PD-L1. B) Comparison of top latent factors predictive of immune phenotypes (inflamed/excluded/desert). C) Comparison of top latent factors with respect to their correlation with tumor mutation burden status of the samples in the cohort D) Survival stratification of the cohort based activation score of top MAUI latent factor (fingerprint) E) Average value of top MAUI latent factors (fingerprints) predictive of response to anti-PD-L1 response groups. F) Comparison of marker combinations in terms of accuracy in predicting anti-PD-L1 response. TMB: tumor mutation burden, CD8_Teff: 12 gene signature for CD8+ effector T-cells, F_TBRS: 4-gene pan-fibroblast TGF-beta response signature, MAUI: top predictive MAUI fingerprints as seen in panel E). Each bar plot represents prediction accuracy (AUC) when using the corresponding subset of variables in an Elastic Net model.

#### Uncovering features of Temozolomide resistance in Glioblastoma Multiforme (GBM)

Temozolomide (TMZ) is a chemotherapeutic agent that is commonly given to patients with Glioblastoma Multiforme (GBM) [48]. TMZ alkylates/methylates DNA, which causes DNA damage, eventually leading to apoptosis. Whereas all patients present intrinsic resistance to TMZ or quickly develop resistance, overall survival is associated with sensitivity to this chemotherapy. Reversal of TMZ-driven alkylates prevents the activation of a DNA damage response and leads to resistance. Tumors with high expression levels of O6-alkylguanine DNA alkyltransferase (AGT), an enzyme which is able to repair the damaging methylation, are more likely to develop resistance to TMZ. In fact, epigenetic silencing through canonical DNA CpG methylation at the promoter of O6-methylguanine-DNA methyltransferase (MGMT), the gene which codes for AGT, is the only available biomarker for TMZ resistance in brain tumors. However, mechanisms to bypass promoter methylation exist, such as MGMT fusion [49] and loss of DNA mismatch repair (MMR), rendering these tumors insensitive to alkylating damage, and thus resistant to TMZ [50].

Thus, MGMT promoter methylation and AGT expression have limited predictive power. Several genome-wide functional screens identified MSI and the Fanconi anemia pathway as critical regulators of response to TMZ. Here, we investigated the ability of MAUI-based multi-omic fingerprints to uncover biomarkers for TMZ resistance directly in GBM biopsies, and to highlight the genes discovered this way. We trained MAUI on 150 samples from the TCGA-GBM cohort [51] for which both mutation and gene expression data were available. Resistant or partially responding tumors were defined as those which experienced tumor progression during the course of treatment with TMZ or in the following six months, while tumors which had six progression free months following the treatment were defined as responders (as has been done for ovarian cancer in [52]) (Fig 6C). Of the 45 patients treated with TMZ, 20 were classified as responding to the treatment, and 25 were classified as non- or partial responders. Using the resulting multi-omic fingerprints, we predicted response/non-response to TMZ using the Lasso with 5-fold cross-validation. Next, we computed the area under the ROC curve (AUC) to quantify each method’s ability to predict TMZ resistance. MAUI fingerprints (AUC=0.88) significantly outperformed both AGT expression (AUC=0.60) and MGMT promoter methylation (AUC=0.68) at this task, but the best performance was obtained by combining MAUI fingerprints with MGMT expression (AUC=0.96) (Fig. 6A). UMAP visualisation based on predictive fingerprints suggest that non-responders sharing the same UMAP space share molecular similarities (Fig. 6B). The difference in AUC is also reflected in statistical significance of progression-free interval differences between the predicted responders and non-responders; stratification based on MAUI fingerprints combined with MGMT expression (log-rank test, p=0.0018, Fig. 6D) outperforms both stratification based on MGMT expression (log-rank test, p=0.23, Fig 6E) and MGMT promoter methylation (log-rank test, p=0.17, Fig. 6F). This demonstrates that MAUI fingerprints capture an additional signal with additional predictive power over MGMT expression or promoter methylation alone.

**Figure 6 :**
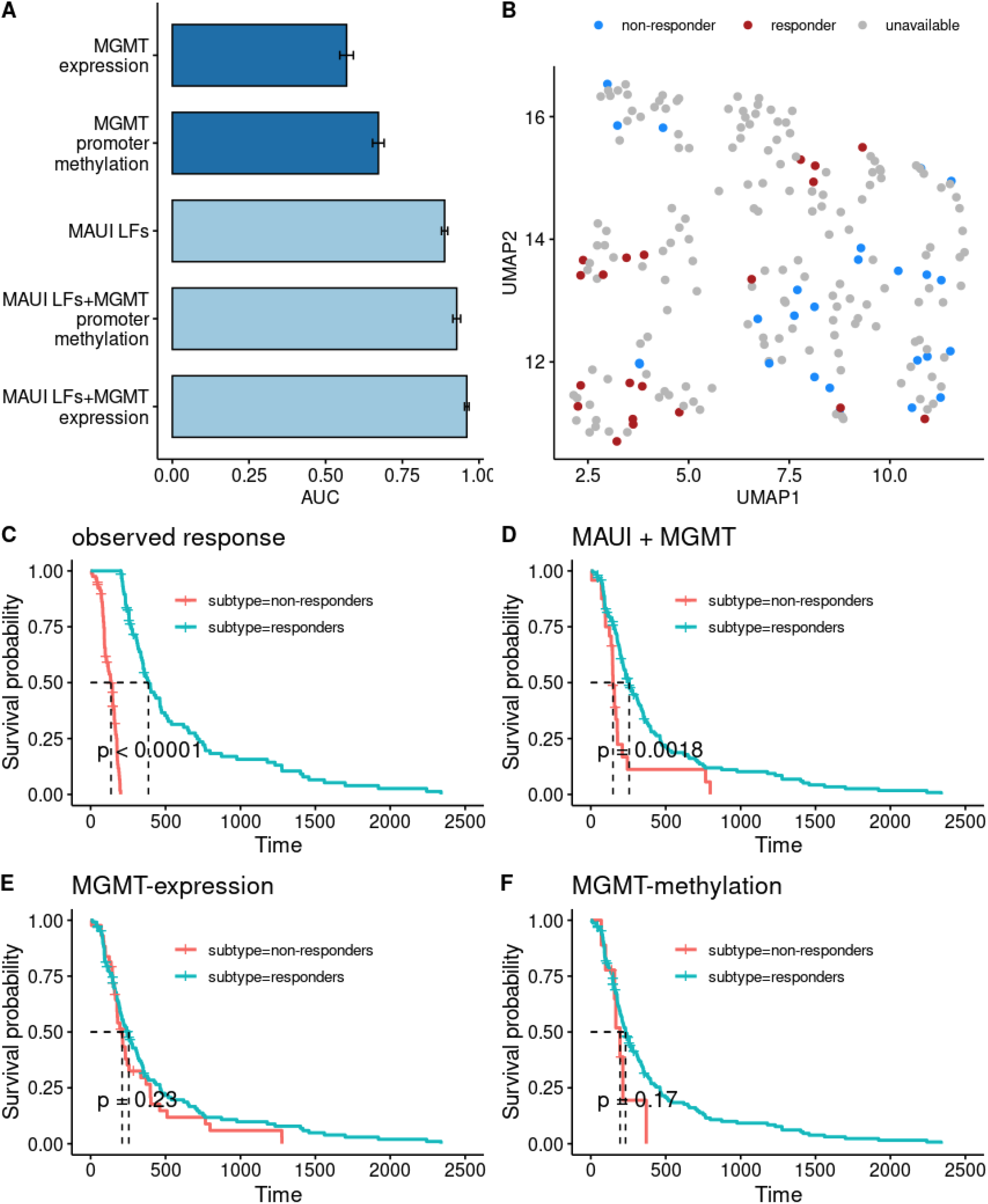
Predicting TMZ resistance in GBM patients. A) Area under the ROC curve (AUC) for 5-fold cross-validation predictions of response to TMZ. MAUI multi-omic fingerprints outperform both MGMT expression and MGMT promoter methylation, and the best performance is achieved by combining MAUI fingerprints with MGMT expression. B) 2D UMAP projection of TCGA-GBM samples based on MAUI multi-omic fingerprints which are predictive of the TMZ response. Responders and non-responders tend to cluster separately. C) Kaplan-Meier curves for progression-free interval for responders and non-responders to TMZ. Response defined as progression-free interval of at least six months, non- or partial-response defined as tumor progression within six months of treatment. D) Kaplan-Meier curves for progression-free intervals for predicted responders and non-responders using MAUI multi-omic fingerprints combined with MGMT expression. E) Kaplan-Meier curves for progression-free intervals for predicted responders and non-responders using MGMT expression. F) Kaplan-Meier curves for progression-free intervals for predicted responders and non-responders using MGMT promoter methylation.

Whereas IDH mutations are strong predictive biomarkers for glioma patients stratification and response to therapy [53], these are not common in GBM. Accordingly, there were only two IDH1 mutants in the TMZ cohort, one of which responded, the other did not. Thus, IDH mutants do not account for MAUI stratification. In order to identify simple biomarkers for this response, we picked the multi-omic fingerprints with nonzero Lasso coefficients, and pulled out the gene coefficients associated with those, multiplied by their lasso coefficients. In the top 100 genes, two features associated with MMR were present (a mutation in MLH1, and the expression levels of PMS2). These two genes have previously been shown to be implicated in TMZ resistance in GBM tumors [54] (Supp. Fig. 4). In addition, many other genes previously shown to be related to TMZ resistance in GBM tumors are flagged (Table 1). The rest of the genes in the list offer a potentially interesting startpoint for further studies into resistance to TMZ, which may be associated with hereto unknown mechanisms of resistance to TMZ, possibly in vivo. Thus, MAUI is able to capture relevant features genetically associated with response to TMZ in brain tumors (e.g. MMR, FA pathways alterations), and may be similarly exploited to investigate response to targeted treatments as mutiomics on these become available.

**Table 1:** Top detected TMZ-resistance related biomarkers that are not members of the MMR pathway, but supported by evidence from the literature.

| Gene | Reference | Remarks |
| --- | --- | --- |
| EEA1 | [55] | Related to increase in overall vesicle production in resistant cell lines |
| FANCI | [56] | Interacts with MMR |
| BIRC3 | [57] | Resistance mediated by inhibition of apoptosis |
| TNFRSF11B | [58] | Involved in methylation regulation |
| C1RL | [59] | Expression predicted resistance in GBM |
| KEAP1 | [60] | Reduction of keap1 by using retinoic acid increases sensitivity to TMZ |
| OTUD4 | [61] | Involved in alkylating damage repair |
| CLK1 | [62] | Also associated with response to cisplatin treatment |
| SOD2 | [63] | Inhibition of SOD2 led to retrieval of drug effect |
| HOXA11 | [64] | When HOXA11 was suppressed in the GBM cell lines, the anticancer effect of temozolomide declined |

## Discussion

In this work, we aimed to demonstrate the applicability and performance of MAUI in its capacity to model the complex interplay between multiple layers of regulatory information in a variety of cancer types for a variety of use cases. This included modelling of clinical parameters, predicting and characterisation of molecular cancer subtypes, prognostic stratification of patients based on survival outcomes, and response or resistance to cancer treatments.

MAUI is a type of autoencoder (a general class of deep learning architectures), specifically a stacked beta-variational auto-encoder. We have utilized MAUI as a tool for joint dimensionality reduction and integration of multi-omics datasets, which is achieved by passing the input multi-omics layers through a “bottleneck layer” of the desired lower dimensionality. Autoencoders consist of an encoder network, which transforms the input data to its latent space representation (found in the bottleneck layer), and a decoder, which reconstructs the input from the latent space representation. The encoder and decoder are trained in tandem, so that the constructed latent space representation can be used to reconstruct the input as faithfully as possible. In doing so, it will capture the essential patterns among the different input features, both within and across different omics modalities. Each neuron in the bottleneck layer represents a single dimension in the latent space, or a single latent factor. Each latent factor is a nonlinear combination of input features, and often corresponds to known cellular molecular structures (i.e. pathways or biological processes). When the multi-omics data comes from a tumor, these patterns have the potential to capture important cancer-related processes: oncogenic mechanisms of action, patient strata markers, predictors of response to therapy, and risk group indicators. Thus, these latent factors represent true multi-omics signatures for biological processes along with technical sources of variation such as batch effects that may have systemic effects on the multi-omics measurements. In this study, we termed the deep learning-based non-linear and combinatorial latent factors “multi-omic fingerprints”.

The multi-omics biomarkers represented by multi-omic fingerprints need not be black-box; rather, they readily lend themselves to interpretation using common bioinformatics toolkits such as gene set analysis [23]. Each fingerprint is a function of a subset of the input genes, and thus each can be said to be affected by a set of genes. The set of genes used by a fingerprint can be compared with gene sets, which are known to be involved in certain biological processes, or to flag certain genes and gene sets as worthy of closer investigation. Hence, multi-omic fingerprints hold the promise of both research tools and complex biomarkers for immediate clinical utility.

We have demonstrated that MAUI multi-omic fingerprints are superior in various settings to latent factors obtained by competing methods. This multi-level information can help researchers view molecular profiles from many different perspectives at the same time. Information contained in MAUI fingerprints can be used in a variety of settings, where some are informative for survival, others might be informative for the common molecular pathways utilized by the subsets of tumors. However, these different perspectives do not necessarily overlap. It is important to stress the relevance of this point in patient stratification in the context of precision medicine. Which patients belong to which stratum depends on the objective of the stratification. For instance, a pair of patients who might belong to the same group based on expected level of overall survival, might be stratified into different groups based on their response to a certain treatment. This is a result of the heterogeneity of how cancer can progress uniquely in each patient and the diversity of the complex molecular patterns that might be associated with different clinical outcomes differently. To further illustrate this argument with an example, we could compare how gastrointestinal cancer patients are stratified based on different outcomes of interest. We observe that patients stratified based on overall survival outcomes can be stratified quite differently if the objective of the stratification was progression-free survival, or if the objective of stratification was to figure out which patients have high MSI status (Suppl. Fig 5A). Although correlations between different groupings exist (Suppl. Fig 5B), each stratification procedure could look like a scrambled version of the allocation of patients by a different stratification procedure.

At its current shape, despite the utility of using MAUI for multi-omics data integration in a non-linear fashion, it also comes with some limitations. As a joint multi-omics integration method, special thought needs to be given to data normalization; different omics data have distinct distributions and dimensionalities, and proper normalization must be applied to avoid giving too much weight to, for instance, the modality with the highest dimensionality, or the least sparse data. It is also not yet optimized to account for the distinct data distributions observed in single-cell omics datasets, that is why although it might work for single-cell multi-omics data integration, other tools such as reviewed here [65] might be more suited for this purpose.

An additional feature that is currently missing in MAUI is the ability to handle missing observations, which can be handled by MOFA [22]. Due to the cost or difficulty of sample acquisition, profiling of all omics layers in every patient is not currently routinely performed. However, on the one hand, the reduction of the cost of data acquisition might overcome this barrier in the future, on the other hand, MAUI is well versed to be applied to large-scale prospective efforts, such as TCGA and similar community-driven approaches.

Another important aspect, which can be thought of as a feature or a limitation depending on the context, stems from the fact that MAUI is a deep learning method. As in other deep learning methods, the latent factors derived from MAUI are not computationally perfectly reproducible. We observed that, even though the exact values of MAUI fingerprints are not identical at every run, we can derive the same conclusions from different runs of the analysis. Moreover, the fingerprints are not necessarily orthogonal to each other or sorted based on total variance explained in the whole dataset as in other linear integration methods such as MCIA, MOFA, or PCA-based methods. From our point of view this is not a limitation, but a feature that gives flexibility in a variety of machine learning tasks where we can discover clinically relevant latent factors for downstream applications. Again, being a deep learning-based method, MAUI requires many samples (we considered cohorts with at least 100 samples in this study) for it to be more useful out of the box, however, it is readily possible to use transfer learning approaches, where the neural networks can be trained in an independent database with many samples and the pre-trained model can be used to further train a relatively small number of samples and make meaningful predictions.

Utilisation of multi-omics data with deep learning algorithms in the context of precision medicine is actually part of a bigger wave of disruptive innovations that are coming about due to the involvement of artificial intelligence approaches in (pre-)clinical research and healthcare in general [66,67]. We believe that deep learning applications of multi-omics profiling along with many other layers of information that can be obtained from personal wearable devices, imaging technologies, and devices that measure the interactions of people with their environment, will become an important tool in not only disease diagnostics, but also continuous health monitoring, disease prevention, prognostics, treatment recommendation, and different levels of clinical/pre-clinical research.

## Methods

### Data

#### TCGA data download and preparation

Omics data from the TCGA consortium was downloaded and prepared for further analysis using TCGAbiolinks R Package [68] for gene expression, mutation, and methylation data. RTCGAToolbox R Package [69] was used to download GISTIC [70] scores for somatic copy number variations for TCGA samples (version date 2016/01/28). The omics platforms included in the analysis are:

1. Gene expression (workflow type: HTSeq - FPKM; data type: Gene expression quantification; data category: Transcriptome profiling).
2. Mutation: Mutation data was downloaded as MAF files.
3. Copy number variation: (data type: GISTIC scores from Broad GDAC Firehose)
4. Methylation: (platform: Illumina Human Methylation 450; data category: DNA Methylation)

#### Clinical Annotation Data

Project specific clinical data was downloaded using TCGAbiolinks [68]. For survival endpoints, we used survival data processed by Liu et al [71].

Tumor purity estimates for TCGA samples were downloaded using the built-in data table *TCGAbiolinks::Tumor*.*purity*. The table includes tumor purity estimates based on immunohistochemistry (IHC), computational estimates by tools such as ABSOLUTE [31], ESTIMATE [32], and LUMP along with a consensus purity estimate [CPE] value derived from the combination of these estimates [30].

#### TCGA subtype annotations

Cancer subtype annotations were downloaded using the *PanCancerAtlas_subtypes* function in the TCGAbiolinks package. The ‘Subtype_Selected’ field was used as the accepted subtype annotation for the corresponding project.

For MSI status prediction, the MSI status annotations by Liu et al [35] were used, where MSI status was quantified using a MSI-Mono-Dinucleotide Assay that examines mono/di-nucleotide repeat loci. MSI annotation data was downloaded using the TCGAbiolinks package.

#### Pan-cancer and pan-organ definitions

TCGA projects were grouped into custom defined projects mentioned in the manuscript.

1. Pan-cancer project includes the following TCGA projects: TCGA-BLCA,TCGA-BRCA,TCGA-CESC,TCGA-COAD,TCGA-READ,TCGA-ESCA,T CGA-HNSC,TCGA-KIRC,TCGA-KIRP,TCGA-LGG,TCGA-LIHC,TCGA-LUAD,TCGA-LUSC,TCGA-PAAD,TCGA-PCPG,TCGA-PRAD,TCGA-SARC,TCGA-STAD,TCGA-T GCT,TCGA-THCA,TCGA-UCEC.
2. Pan-gastrointestinal project includes: TCGA-ESCA, TCGA-STAD, TCGA-COAD, TCGA-READ.
3. Non-small-cell Lung Cancer project includes: TCGA-LUAD, TCGA-LUSC.

#### Omics data preparation for multi-omics integration

##### Input features

For training MAUI, we collated a gene set consisting of 7679 genes by taking a union of all the cancer hallmark gene sets from the MSIGDB database [72] and tumor microenvironment related gene set annotations from the xCell R package [73].

1. Gene expression: Genes were sorted by most variable FPKM counts and top 2500 genes were kept for further analysis.
2. Copy number variation: Gene-level copy number scores from GISTIC [70] were used. Top 2500 genes with most variable GISTIC scores were kept for further analysis.
3. Mutation: Mutations of any variant classes were extracted from the MAF files for each project. For each gene, the number of mutations were counted. The genes were sorted by the number of samples in which the gene is mutated. Top 2500 genes were kept for further analysis.
4. Methylation: ∼450,000 CpG methylation sites were firstly filtered for those that overlap the promoters of the selected set of collated genes. The remaining CpG sites were sorted by variance of methylation scores across the analyzed cohort of samples. Top 2500 CpG sites were kept for further analysis.

##### Input Samples

For each of the omics data types, the normal samples (TCGA sample type codes: ‘NB’, ‘NBC’, ‘NEBV’, ‘NT’, ‘NBM’) were excluded to only consider tumor samples in the analysis. The remaining samples were grouped by patient barcodes (bcr_patient_barcode). For patients with more than one sample per data type, the first sample by alphabetical order was kept. Finally, patients that lack at least one sample per data type were excluded and the obtained data matrices were individually scaled by column. The input for multi-omics integration tools is eventually 4 data matrices (one for each data type), each containing omics profiles for the same set of patients.

#### Immunotherapy Response in Metastatic Urothelial Carcinoma Dataset

Gene expression measurements of a cohort (N = 348) of Metastatic Urothelial Carcinoma pre-treatment with anti-PD-L1 was downloaded from the IMvigor210CoreBiologies R Package (version 0.1.13) as described in [46]. The raw gene counts were normalized by the sample-wise scaling factors as provided in the data package and further log-transformed. Genes were further filtered to keep only those that are members of the cancer hallmark gene sets from the MSIGDB database and the xCell gene set annotations. Top most variable 5000 genes were selected as input features. The annotations for treatment response, tumor mutation burden, and immune phenotypes were all used as provided in the IMvigor210CoreBiologies R package.

### Multi-omics integration tool settings

#### MAUI

maui-tools version 0.1.93 [23] was run with 1000 hidden units, for 500 epochs, looking for 100 latent factors. Feature contributions to each latent factor were extracted using the *get_neural_weight_product* function. For unsupervised clustering tasks, MAUI latent factors that are highly correlated were filtered prior to the clustering task using the *findCorrelation* from the caret R package [74]. The correlation cut-off was set to the value that corresponds to the 99th percentile of the pairwise correlation distribution of the corresponding matrix of latent factors.

#### MOFA

MOFA2 version 1.0.0 [22] was run with the default settings except for the number of factors. The number of factors was set to a quarter of the number of input samples with an upper limit of 100 factors (if the number of samples is larger than 400). Uninformative factors were automatically dropped by the algorithm.

#### PCA

The *prcomp* function in the built-in *stats* library in R [75] was used to compute 100 principal components.

#### MCIA

omicade4 (v.1.30.0) R package [21] was used to train the Multi-Co-Inertia Analysis algorithm. The default settings were used searching for 20 factors for each project. Our initial attempt to search for 100 factors failed, because the algorithm didn’t converge for that many factors. The number of searched factors was decreased to 20, which worked for all experiments.

### Machine learning tasks

#### Classification

Random Forest [76] models were built using the ranger R package [77] with a 60/40 train/test data partitioning with default settings, but looking for 1000 trees unless otherwise stated.

For Elastic Net models, *caret* R package [74] was utilized to run a 5-fold cross-validation to optimize hyperparameter settings of Elastic Net by glmnet R package [39] on the training dataset (60% of the data). Area under the ROC curve (AUC) and the corresponding confidence intervals were computed using pROC R package [78].

#### Detecting Latent Factors associated with discrete variables

In order to quantify the predictive power of latent factors or individual omics features for discrete variables such as clinical parameters (e.g. tumor stage) and molecular subtypes (e.g. MSI status), for each subgroup of patients corresponding to a subgroup of a given discrete variable, a one-vs-all Wilcoxon rank-sum test was performed (alternative = greater) to detect latent factors (or omics features) that are enriched for the subgroup of patients. The p-values obtained from the statistical test were further adjusted for multiple testing using the Benjamini-Hochberg method as implemented in the stats R package [75].

#### Ranking input features by contribution to latent factors

The individual contribution of each input omics feature to each latent factor was computed and ranked according to the absolute neural path weight products, which is defined as the product of weights along the path from from input features to latent factors, using the *neural_path_weight_product* function of MAUI package [23].

#### Building decision trees from omics features for non-small cell lung cancer subtypes

Top 10 omics features contributing to each of top 5 latent factors predictive of the dependent variable (e.g. lung cancer subtypes) were detected based on neural path weight product (nwp)-rankings of omics features per each LF. The candidate omics features were further used to build an initial conditional inference tree on 60% of the samples. Top most important features from this model were extracted to build a final model on the 60% of the samples. The final decision tree classification performance was evaluated on the remaining 40% of the samples. Conditional inference (decision) tree was built using the partykit R package [79,80] and visualised using the treeheatr R package [81].

#### anti-PD-L1 (immunotherapy) response prediction

MAUI was trained on the full cohort of 348 samples with gene expression measurements. For patients with tumor mutation burden (TMB) measurements (N = 216), the dataset was split into training/testing in a 60/40 split. An initial Elastic Net model (5-fold cross-validation with 5 repetitions using down-sampling to account for class imbalance) was built to detect 5 MAUI LFs that are predictive of response to therapy.

These top 5 LFs were used in combination with published response markers in the study by Mariathasan et al [46], which included TMB, a 12-gene signature for CD8+ effector T-cells, and a 4-gene signature for pan-fibroblast TGF-beta response. Combinations of the different sets of markers were used to build Elastic Net models (5-fold cross-validation with 5 repetitions using down-sampling to account for class imbalance) to predict the responders to anti-PD-L1 therapy. The models were evaluated on the test dataset using the area under the ROC curve (AUC) metric.

‐ CD8+ effector T-cell (CD8_Teff) signature genes: CD8A, GZMA, GZMB, IFNG, CXCL9, CXCL10, PRF1, TBX21
‐ pan-fibroblast TGF-beta response (F-TBRS) genes: ACTA2, COL4A1, TAGLN, SH3PXD2A

### Survival analysis

We used Progression Free Interval (PFI) or Overall Survival (OS) annotations from Liu et al [71]. Harrell’s C-index as a measure of survival outcome prediction accuracy was computed by fitting a Random Survival Forest using the randomForestSRC R package [82] and computing the average error per each grown tree (1000 trees). We followed a two-step fitting procedure, where we initially built a random survival forest on the training data using all latent factors, from which we picked the top 10 latent factors in terms of variable importance for predicting survival in the training data. A second model was built using the top 10 survival-associated latent factors. This final model was evaluated on the testing data by computing the C-index values. survival R package [83] was used to build survival objects, fitting Cox Proportional Hazards models, and computing corresponding hazard ratios. Log-rank tests and covariate-adjusted Kaplan-Meier curves were computed using the survminer R package [84].

### Modelling tumor purity estimates

Elastic Net regression models (using repeated five-fold cross-validation) were built on 60% of the samples for each TCGA cohort, for which the tumor purity estimates were available (14 TCGA cancer types out of the 21 studied cancer types), where the input features were MAUI latent factors and the dependent variable was the reported tumor purity estimates by IHC or CPE. The models were evaluated on the remaining 40% of the samples with respect to the Pearson correlation between the predicted tumor purity values and the reported tumor purity estimates (IHC or CPE).

### Modelling TMZ resistance in GBM

Response to TMZ was defined as no tumor progression for six months following the last treatment. Partial and non-responders are defined as patients who experienced tumor progression in that same six month period. Prediction of TMZ response was done using the Lasso (l1-regularized logistic regression) tuned using 5-fold cross-validation, with each sample predicted when it was not in the training set for a full set of out-of-training-set predictions. The input to the Lasso was the set of latent factors learned by MAUI, and the output response/non-response vector was as defined here. The ROC was computed and the area under the curve calculated to quantify the prediction accuracy. Error bars were calculated by bootstrapping the entire procedure. AUCs for MGMT expression and promoter methylation were calculated likewise, using a simple logistic regression without regularization.

### Clustering

Clustering experiments that were carried out to evaluate the stability and homogeneity of clusters were done using the clValid R package [28]. Unless otherwise stated, the k-means algorithm implemented in the stats R package [75] was used to cluster samples with 1000 random restarts and a maximum 30 iterations.

## Data visualisation

The plots in the manuscript were generated using ggplot2 [85], ggpubr [86] and cowplot [87] libraries. Heatmaps were plotted using pheatmap R package [88]. tSNE dimensions were calculated using the Rtsne R package [27]. Decision trees were computed using the partykit R package [79] and visualised using the treeheatr R package [81].

## Data and Code availability

All data used in this study are publicly available. The code that was used for downloading, preparing, and processing the publicly available data, training of the multi-omics integration tools, and generating the figures in the manuscript can be found at https://github.com/BIMSBbioinfo/uyar_et_al_multiomics_deeplearning.

## Acknowledgements

We thank the members of the Akalin Lab for their valuable feedback to the project and Ricardo Wurmus, Dan Munteanu, and Madalin Patrascu for maintaining the group servers that provided us a reliable and efficient computational environment. The G. Gargiulo lab acknowledges funding from MDC, Helmholtz (VH-NG-1153), ERC (714922).

The results of this study are in whole or part based upon data generated by the TCGA Research Network: https://www.cancer.gov/tcga.

## Author Contributions

AA and JR conceived the study. BU and JR did the formal analysis. BU, JR, VF, and AA wrote the paper with contributions from GG. All authors read, edited, and confirmed the final manuscript. AA acquired funding and supervised the project.

## Manuscript Figures

**Supplementary Figure 1 :**
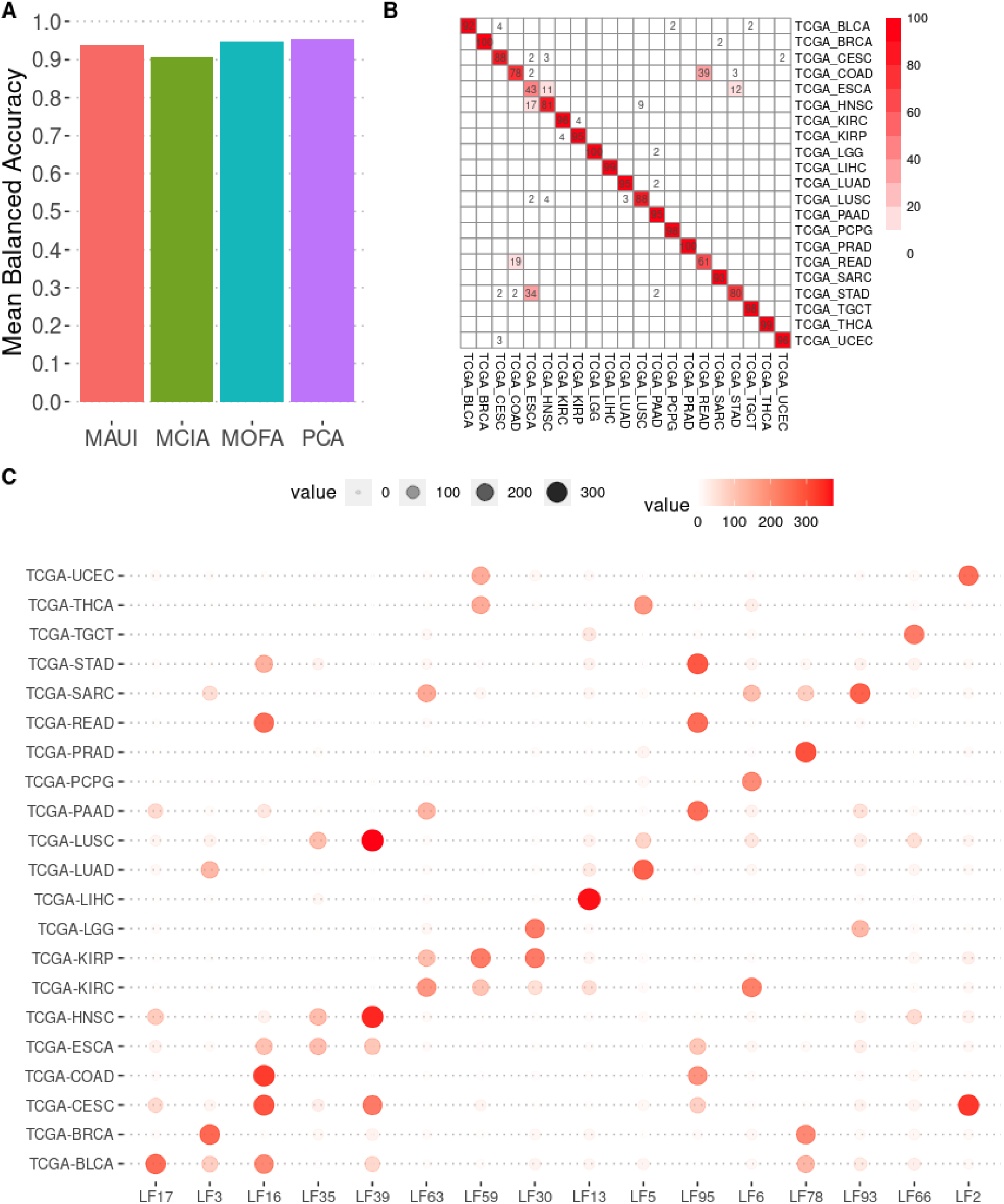
A) Predictive power of latent factors in distinguishing cancer types comparing MAUI with PCA/MOFA/MCIA. Mean Balanced Accuracy computed on the test dataset (40% of held-out data) trained on 60% of the dataset using an Elastic Net multi-class model. B) Confusion matrix of predicted versus true labels of samples’ cancer types. The color of each cell represents the percentage of samples from the cancer type in the rows that are classified into the cancer types on the column. Cells that deviate from the red color represent a relatively lower level of classification accuracy. C) Top MAUI latent factors (multi-omic fingerprints) detected per cancer type. Size/color/opacity of the points represent the average neural activation value of the MAUI fingerprint for the corresponding type of tumor samples.

**Supplementary Figure 2 :**
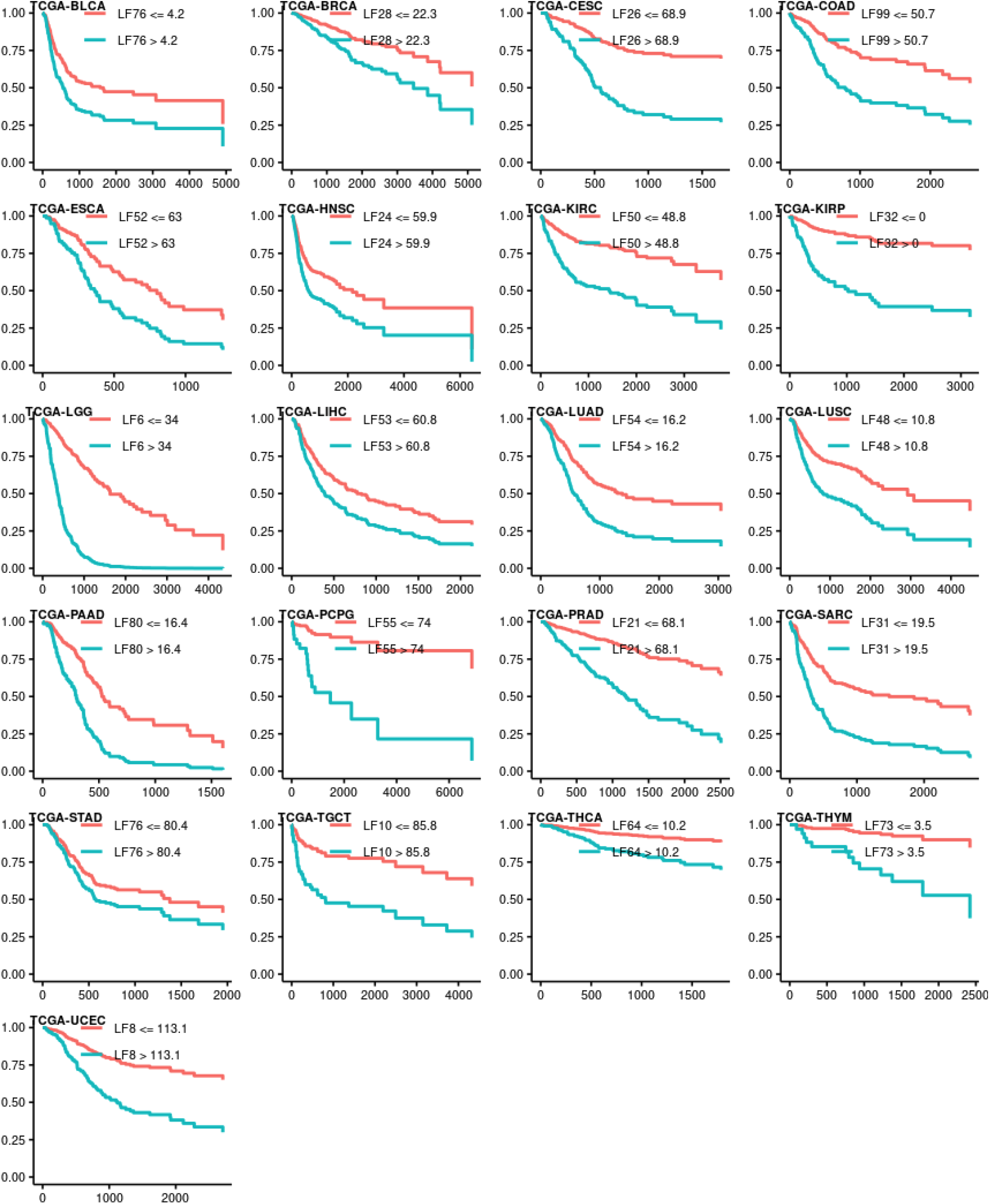
Kaplan-Meier survival curves adjusted for clinical factors (tumor stage + age + gender) stratified by the neural activation values of top survival-predictive multi-omic fingerprints per cancer.

**Supplementary Figure 3 :**
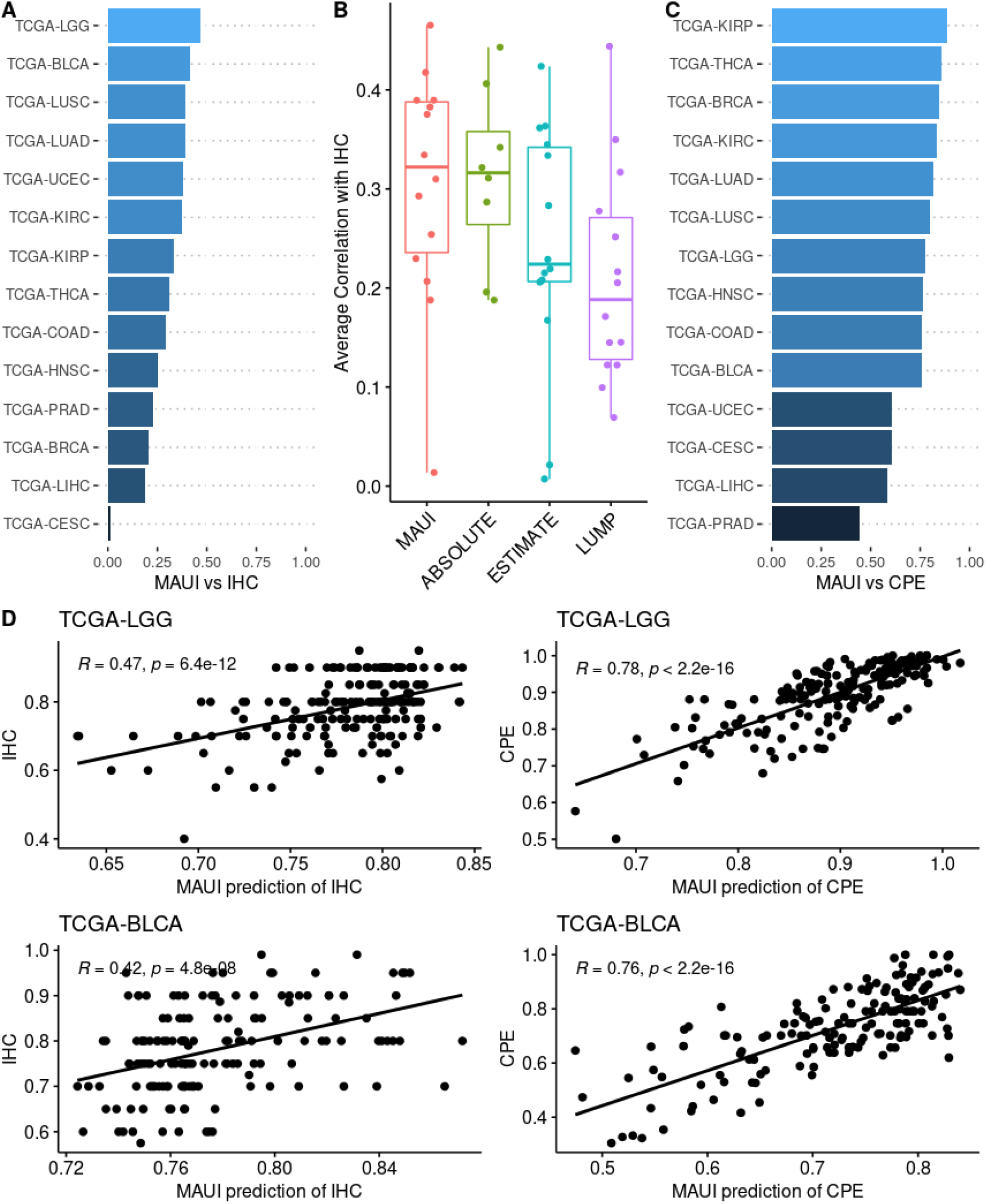
Predicting tumor purity estimates using MAUI latent factors. A) Correlation of the MAUI predictions of IHC versus the annotated IHC values (immunohistochemistry-based tumor purity estimation). B) Comparison of average correlations between computational estimates of IHC values by different tools. Each point represents the value obtained for each of the TCGA cohorts seen in panel A. ABSOLUTE tool lacks estimates for 6 out of 14 cancer types. C) Correlation of the MAUI predictions of CPE versus the annotated CPE levels (CPE stands for consensus purity estimated by computational methods: ABSOLUTE, ESTIMATE, and LUMP). D) Scatterplots of MAUI predictions versus the annotated IHC and CPE values for the top two best predicted cancer types with respect to annotated IHC values.

**Supplementary Figure 4 :**
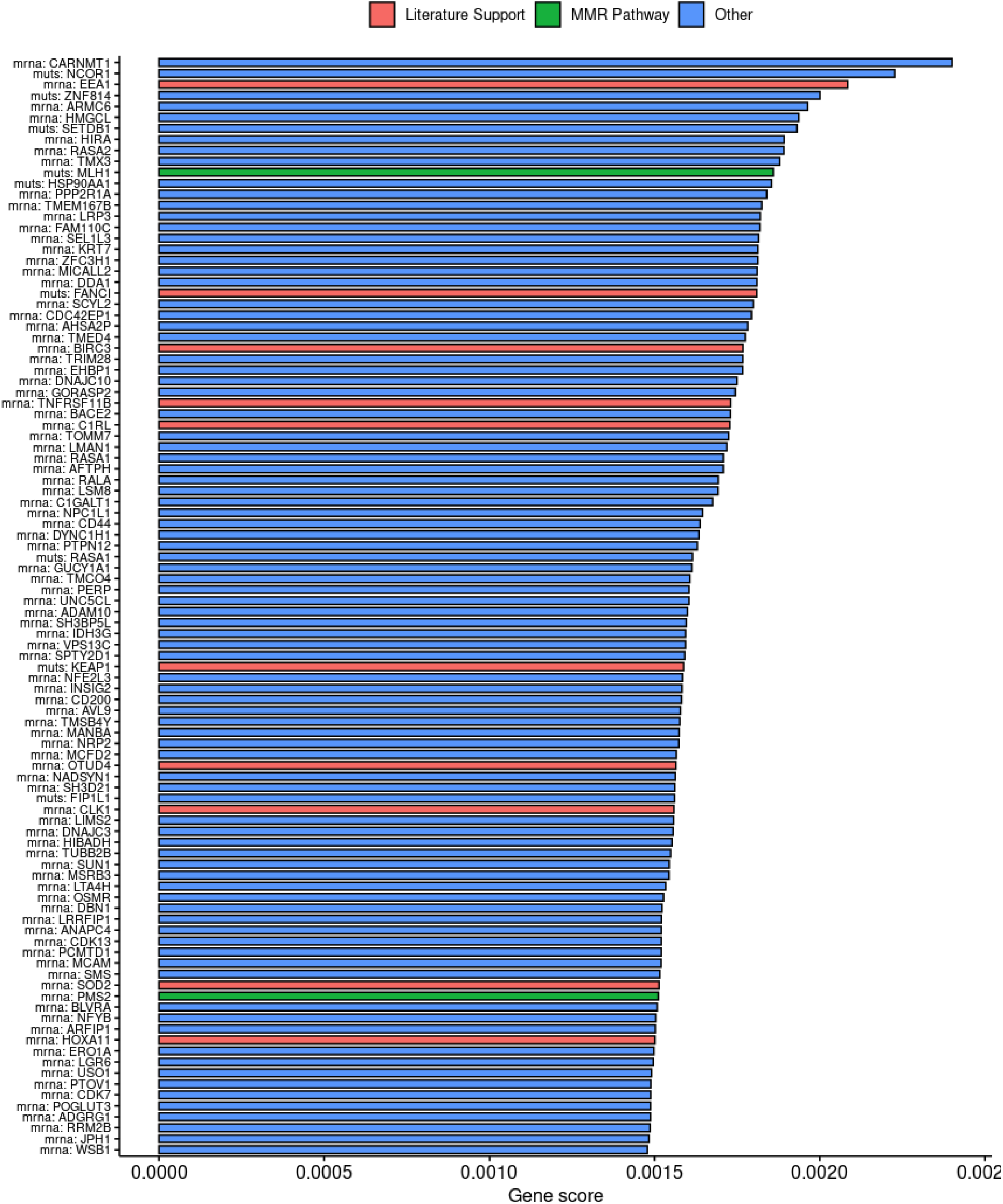
Top 100 genes sorted by importance in predicting TMZ resistance in GBM patients.

**Supplementary Figure 5 :**
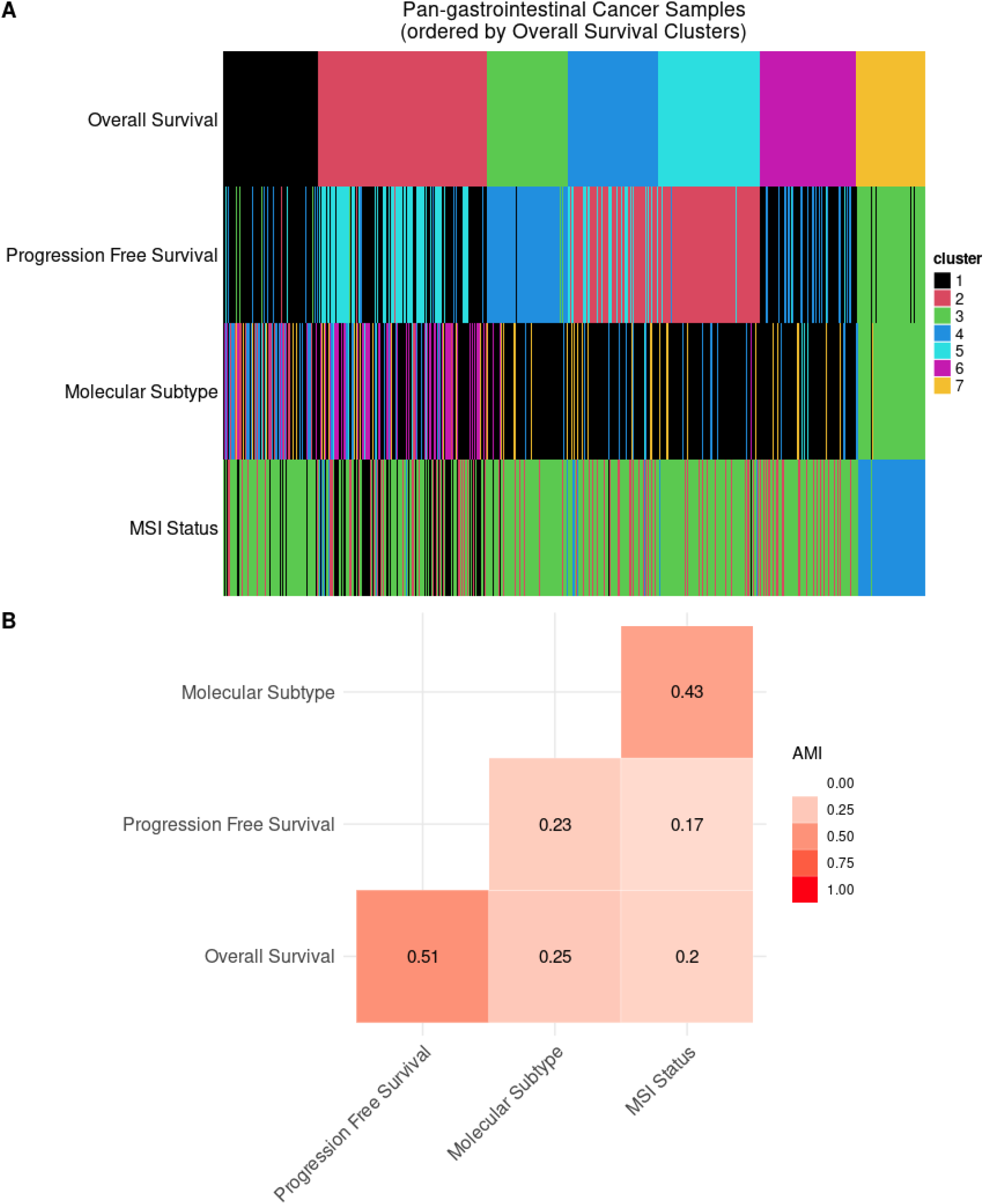
Demonstration of how cancer subtypes are context/objective dependent. A) Heatmap of patient cluster memberships based on different objectives: optimum stratification based on overall survival or progression free survival, molecular homogeneity (TCGA pan-GI subtypes), or micro-satellite instability (MSI) status. Columns are ordered first by membership in the optimum Overall Survival cluster membership. B) Pairwise Adjusted Mutual Information (AMI) of cluster labels obtained via different patient stratification methods. The AMI value ranges from 0 to 1 and the higher the value, the more overlap between patient partitions.

